# Mammalian Cell Entry domains are required for bile resistance and virulence in *Salmonella*

**DOI:** 10.1101/263871

**Authors:** Georgia L. Isom, Jessica L. Rooke, Camila A. Antunes, Emma Sheehan, Timothy J. Wells, Christopher Icke, Adam F. Cunningham, Jeffrey A. Cole, Ian R. Henderson, Amanda E. Rossiter

**Affiliations:** Institute of Microbiology and Infection, University of Birmingham, UK; Microbiology Division, Friedrich-Alexander University Erlangen, Nuremberg, Erlangen, Germany

## Abstract

MCE domains were first reported in *Mycobacteria* as having a role in <u>M</u>ammalian <u>C</u>ell <u>E</u>ntry, with subsequent studies showing their importance during infection. Here, we have examined the function of MCE proteins in *Salmonella* Typhimurium during mammalian infection. We report that MCE proteins are required for *Salmonella* virulence, but that this is not related to decreased adherence, entry or survival in mammalian cells. Instead, we reveal that MCE proteins are required for *Salmonella* bile resistance, in particular to withstand bile salts such as cholate and deoxycholate. Based on our previous work in *Escherichia coli*, and other studies that have reported roles for MCE proteins in membrane biogenesis, we propose that *Salmonella* lacking MCE domains have a defective outer membrane that results in bile sensitivity and decreased virulence *in vivo*. These results suggest that MCE domains mediate fundamental aspects of bacterial membrane physiology as opposed to a proposed direct role in mammalian cell entry, explaining their conservation across both pathogenic and non-pathogenic bacteria.

## Introduction

Mammalian cell entry (MCE) domains are protein domains that were first identified in *Mycobacterium tuberculosis,* where insertion of a gene from *M. tuberculosis* into non-pathogenic *Escherichia coli (E. coli)* was shown to allow the bacterium to enter and survive in mammalian cells^1^. Because of this acquired ability, this gene was named *mce1A,* for mammalian cell entry. MCE domains are widespread across diderm bacteria^2^ and have been most thoroughly studied in Actinobacteria, especially in *Mycobacteria*.

In the last few years, research in MCE domains has expanded to include a focus on Proteobacteria, reporting roles in lipid trafficking and membrane biogenesis^2,3,4^ Despite the well-established role of Actinobacterial MCE domain-containing proteins (“MCE proteins”) in mammalian infection^5,6,7^, research into Proteobacterial MCE domains during mammalian infection is sparse.

Here, we aimed to investigate for the role of MCE domains in Proteobacterial mammalian pathogenicity, *in vivo.* For this purpose, we used the clinically important model organism *Salmonella.* It is primarily a pathogen of the gastrointestinal tract, but serovars such as *Salmonella enterica* subsp. *enterica* serovar Typhimurium *(S.* Typhimurium) and *Salmonella enterica* subsp. *enterica* serovar Typhi *(S.* Typhi) can cause systemic infection in mice and humans, respectively, resulting in spread to organs such as the liver, gall bladder and spleen. *S.* Typhimurium murine pathogenesis is well understood and provides an infection model for both *S.* Typhimurium (causing gastroenteritis) and *S.* Typhi (causing typhoid fever) in humans.

The *Salmonella* genome encodes three MCE proteins: MlaD, PqiB and YebT. In *E. coli,* all three of these proteins have been implicated in phospholipid transport important for maintenance of the outer membrane^3,4^, and homologues of these proteins are important for infection *in vitro.* A homologue of MlaD found in *Shigella flexneri* is required for intercellular spreading^8^, and a YebT homologue in *Vibrio parahaemolyticus,* is reported to be an adhesin involved in early infection that binds to mammalian cells^9,10^. Furthermore, *in vivo* studies in insects and nematodes^9,11^ support a role for Proteobacterial MCE proteins during pathogenesis.

The work presented here shows that MCE proteins are required for *Salmonella* virulence during systemic infection in mice, a result potentially applicable to other Proteobacterial mammalian pathogens. Despite the term ‘mammalian cell entry’ domain, we show that this infection phenotype is not dependent on mammalian cell entry or survival within the mammalian cell, but that these proteins are important for bile resistance, particularly to bile salts such as cholate and deoxycholate.

## Results

### MCE proteins are required for systemic infection of *Salmonella* Typhimurium

To determine the importance of MCE proteins during murine infection, a triple deletion mutant of *mlaD, pqiAB* and *yebST* (MPY mutant) was constructed in *S.* Typhimurium SL1344 (WT) using a combination of gene doctoring^12^ and P22 transduction. Prior to murine infection, both the WT and MPY mutant were grown in lysogeny broth (LB) to check for general growth defects. No differences were observed (Supplementary Figure 1).

Next, 6 mice per strain were infected via intraperitoneal injection with 1000 bacteria. After 3 days, the bacterial burdens in the liver and spleen were measured by plating organ extracts onto LB agar plates. The number of colony forming units (CFUs) in the livers and spleens from mice infected with the MPY mutant were 26 times and 34 times lower than in the WT (Figure 1). We also showed that a single *mlaD* mutant and a double *pqiAB yebST* mutant were also attenuated, but to a lesser extent than the triple mutant, indicating that all three MCE proteins contribute to the observed attenuation (Supplementary Figure 2).

**Figure 1.**
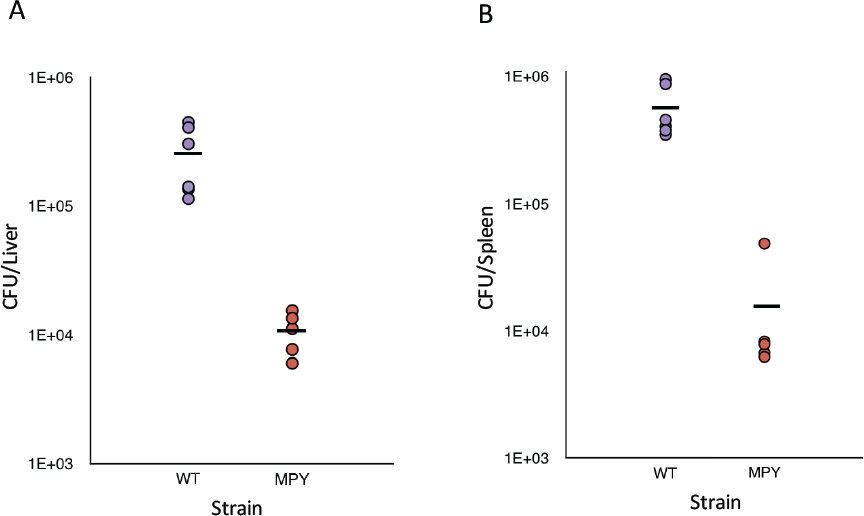
Bacterial burden of *Salmonella* Typhimurium WT and *mlaD, pqiAB, yebST* mutant (MPY) in C57BL/6 mice. Six mice were infected with the strains for 3 days via intraperitoneal injection. The CFUs/organ were counted in the liver **(A)** and spleen **(B)**. Each point represents the number of CFUs/organ from a single mouse, and the line indicates the mean number of CFUs. Significant differences between the WT and MPY
mutant were tested using a Mann-Whitney U test - liver: p=0.004, spleen: p=0.004).

### MCE proteins are not required for serum resistance

To determine at which stage of infection MCE proteins are required, we tested the abilities of the WT and mutant strains to resist complement-mediated killing in serum, which is important for establishing infection *in vivo. Salmonella* cultures were mixed at a 1:9 ratio with serum obtained from healthy humans, and samples were taken after 0, 45, 90 and 180 minutes and plated onto LB agar. For comparison, a *S.* Typhimurium SL1344 strain lacking O-antigen and known to be serum-sensitive^13^ was also tested, and this strain exhibited a clear drop in CFUs after 45 minutes. In contrast, the WT and MCE triple mutant were equally resistant (Figure 2). These results indicate that MlaD, PqiB and YebT are not required for *S.* Typhimurium resistance to serum and that the attenuation observed *in vivo* is not related to the ability of mutant to withstand complement-mediated killing.

**Figure 2.**
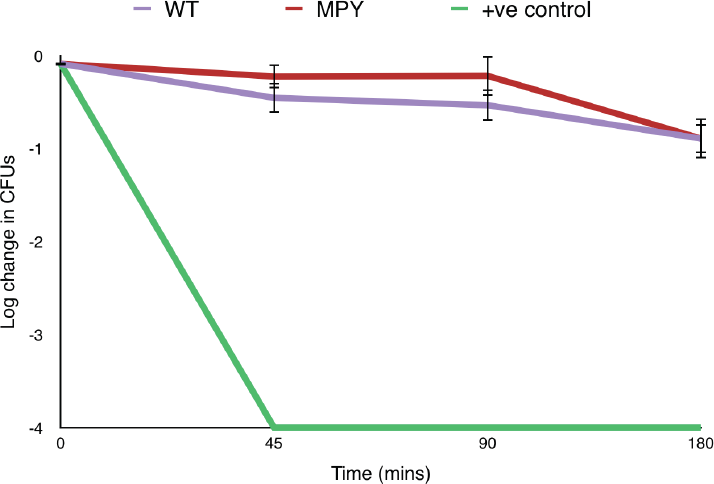
Bacterial survival of the *S*. Typhmurium WT and MPYmutant in the presence of healthy serum. The positive control was a strain of S. Typhmurium SL1344 lacking O-antigen. Each point is the mean of 3 biological replicates and the error bars represent the standard deviation.

### MCE proteins are not required for adhesion or invasion of non-phagocytic mammalian cells

A key step of *Salmonella* infection is its self-promoted uptake into non-phagocytic cells^14^. Therefore, we wanted to determine whether MCE proteins are important for this process, by comparing the ability of the WT and the triple MCE mutant strain to adhere and invade HeLa cells. The bacteria were added to HeLa cells in a ratio of 10: 1 and left for either 30 minutes to measure adherence, or for an additional 2 hours with gentamicin treatment to measure invasion. At each time point the HeLa cells were lysed, bacteria were plated, and the CFUs were calculated. For both adherence and invasion, there were no significant differences between the WT and the MPY mutant (Figure 3). Therefore, it is unlikely that the attenuation observed during infection in mice was due to a decrease in mammalian cell adherence or entry.

**Figure 3.**
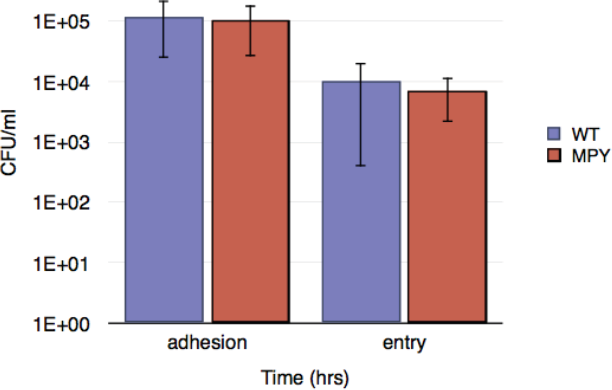
Adhesion and invasion of HeLa cells with S. Typhimurium WT and MPY mutant. For adhesion, the bacteria were left to adhere to HeLa cells for 30 minutes. For the 2-hour time point, the bacteria were given an additional 2 hours in the presence of gentamycin to test for invasion only. The bars are a mean of 3 biological replicates and
the error bars represent the standard deviation. Significance was determined using a student's t-test: adhesion: p=0.79, entry: p=0.47).

### MCE domains are not required for survival in macrophages

Macrophages are an important part of the host immune response, responsible for phagocytosis and killing of invading pathogens. *Salmonella* have evolved to resist killing in macrophages and instead use them as a vehicle for systemic infection^15^. Therefore, we wanted to test whether the MPY mutant strain was less able to survive in macrophages.

We infected bone marrow derived macrophages with the WT and MPY mutant bacteria for 2, 5 and 24 hours, and the number of viable bacteria was determined by counting CFUs. There were no significant differences between the ability of the WT and MPY mutant to survive in bone marrow-derived macrophages (Figure 4). Therefore, we concluded that the attenuation observed *in vivo* was not related to survival in macrophages.

**Figure 4.**
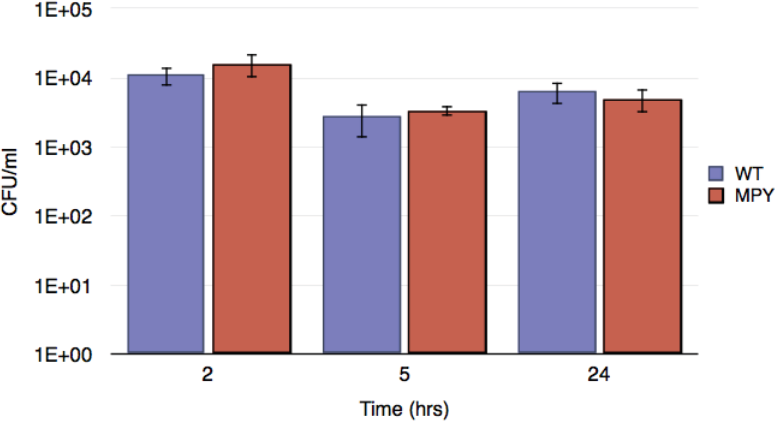
Survival of the WT and MPY mutant in bone marrow derived macrophages after 2, 5 and 24 hours of invasion. The bars are a mean of 3 biological replicates and the error bars represent the standard deviation. Significance was determined using a student's t-test: 2 hrs: p=0.30, 5 hrs: p=0.61, 24 hrs: p=0.44).

### MCE proteins are required for bile resistance

During systemic infection, *S.* Typhimurium must survive in bile to colonise the gall bladder and be subsequently released into the intestinal lumen for shedding and spreading between hosts^16^. Therefore, *Salmonella* can withstand toxic substances in bile, such as bile salts, and can also use compounds, especially phospholipids, in bile as nutrient sources^17^. To determine whether a mutant lacking MCE proteins was less able to survive in bile, the WT and MPY mutant strains were plated onto LB agar containing bile, bile salts, cholate or deoxycholate, or on minimal agar containing phosphatidylcholine or phosphatidylglycerol as a sole carbon source. The MPY mutant was much less able to grow in the presence of bile than the WT (∼100 times, Figure 5). A similar growth defect was observed in the presence of bile salts, cholate or deoxycholate. In contrast, no growth differences were observed between the WT and MPY mutant on phospholipids. This suggests that the impaired growth of the MPY mutant on bile is due to sensitivity to bile salts rather than a defect in nutrient utilisation. This is supported by the fact that the growth defect worsens with increasing bile concentration (Supplementary Figure 3). The single *mlaD, pqiAB* and *yebST* mutants and double *pqiAB yebST* mutants were also more sensitive in the presence of bile (Supplementary Figure 4). Overall, these results indicate that the attenuation observed *in vivo* was related to the inability of the MCE mutants to survive in bile, due to increased susceptibility to bile salts.

**Figure 5.**
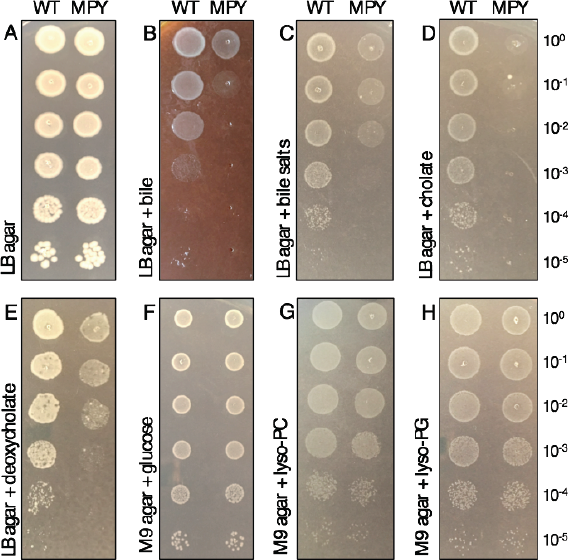
Logarithmic dilutions of the WT and MPY mutant on A) LB agar only, B) LB agar supplemented with 15% bile, C) LB agar supplemented with 5% bile salts, D) LB agar supplemented with 5% cholate, E) LB agar supplemented with % deoxycholate, F) M9 agar supplemented with 0.4% glucose, G) M9 agar supplemented with 100 µg/ml phosphatidylcholine and H) M9 agar supplemented with 100 µg/ml phosphatidylglycerol.

## Discussion

Here, we use *Salmonella* Typhimurium to show that MCE proteins in Gram-negative bacteria are important for mammalian infection *in vivo.* These findings are in agreement with studies in other organisms, such as *Mycobacterium tuberculosis*^5,6,7^, suggesting that MCE proteins are important for virulence across pathogenic diderm bacteria. Investigation into this attenuation phenotype revealed that it is unlikely to be due to serum sensitivity or adhesion, invasion, or survival within mammalian cells. Instead, we show that MCE proteins in *Salmonella* are important for survival in bile.

As MCE domains were originally described to have a direct involvement in mammalian cell entry^1^, the results presented here might appear counter-intuitive. However, few studies have actually shown that mutants lacking MCE proteins display impaired ability to enter mammalian cells. The Mce1A protein found in *M. tuberculosis* was the original protein shown to aid *E. coli* entry and survival into macrophages^1^. Whilst beads coated with Mce1A or bacteria overexpressing Mce1A show increased entry into HeLa cells^1,18,19,20^, *mcel* mutants do not show a decreased ability to enter and survive in mammalian cells^5,7^. Due to their lipid-binding properties^2^, it is possible that isolated MCE proteins may directly bind to the surface of mammalian cells. However, in native conditions, these proteins locate to the cytoplasmic membrane of Gram-negative bacteria and organelles^2,4,8,21^, making it unlikely that they directly interact with mammalian cells to induce cell entry. Therefore, we propose that whilst MCE proteins could indirectly affect mammalian cell entry, this is not their primary function. This is supported by the more recent literature implicating MCE proteins in lipid trafficking for outer membrane biogenesis^2,3,4^ and the fact that MCE domains are present in a wide range of non-pathogenic bacterial and eukaryotic species^2^.

Instead, we have shown that MCE proteins in *S.* Typhimurium play an important role in bile resistance *in vivo,* in particular resistance to bile salts such as cholate and deoxycholate. Bile salts are detergents that are designed to emulsify fats in the human gut. However, they can also disrupt bacterial membranes^22^. Given our previous data linking MCE proteins to outer membrane biogenesis^2^, and the fact that mutants in *E. coli* are sensitive to detergents^2,3,4^, it is likely that the phenotypes observed here are due to a defective membrane of the *Salmonella* MCE mutants. This is consistent with the fact that the membranes of *mce* mutants in *Mycobacteria*^23^ and *Streptomyces^24^* are also perturbed. However, it is important to mention that despite the binding of MlaD, PqiB and YebT to phospholipids, and the proposed role of MCE proteins in lipid uptake in Actinobacteria, the *Salmonella* MPY mutant can still import and utilise phospholipids. This suggests that MlaD, PqiB and YebT are not involved in, or are not solely responsible for, phospholipid uptake from the extracellular environment.

In summary, we provide further evidence that MCE domains are not directly involved in mammalian cell entry, but are instead required for the maintenance of the bacterial outer membrane. As a consequence, MCE proteins in *Salmonella* are required for murine infection, particularly to resist the detergent effects of bile salts. However, due to the conservation of these proteins across both pathogenic and non-pathogenic bacteria, we conclude that MCE proteins are important for virulence but have not evolved specifically for this purpose.

## Methods

### Strains, media and growth conditions

*Salmonella* Typihurium SL1344 was used as the parent strain. Bacteria were grown in lysogeny broth (LB) or on LB agar plates (LB supplemented with 1.5% nutrient agar) and grown at 37°C. To construct deletions in *mlaD, pqiAB* and *yebST,* the genes were replaced by a kanamycin resistance cassette using gene doctoring, as previously described^12^. The kanamycin cassette was removed using the vector pCP20. Double and triple mutants were constructed by transfer of the kanamycin resistance cassette gene replacements into the same background, with using P22 transduction.

For growth curves, strains were inoculated into 50 ml of LB medium at a starting OD_600_ of 0.05. Time points were taken once an hour for 8 hours. For dilution plates, cultures were adjusted to an OD_600_ of 1 and diluted down to 10^−5^ in a microtitre plate. A multichannel pipette was used to transfer 2 µl of the dilutions for each strain to LB or M9 agar plates supplemented with the desired chemicals.

### In vivo *infection of mice*

For *in vivo* infection a culture of each strain was grown to an OD_600_ of 1. The cells were centrifuged and washed twice with PBS. After a final re-suspension in PBS, the concentration of cells was adjusted to a concentration of 5000 cells/ml. For injection, each mouse was injected with 200 µl of bacteria (1000 bacteria). After 3 days of infection, the mice were culled and the livers and spleens were isolated. The organs were mashed through a cell strainer using a sterile syringe and rinsed through with 5 ml of PBS into a 50 ml tube. The mixture was diluted down to 10^−3^ and plated onto LB agar plates and incubated overnight at 37°C.

### Serum bactericidal assay

For each strain, 1 ml of culture at OD_600_ of 1 was centrifuged, washed twice and resuspended in PBS to make the strain stock inoculum. The stock inoculum was diluted 1:10 and 10 µl was added to 90 µl serum and incubated for 45, 90 and 180 min at 37°C. At each time point a sample was collected and diluted down to 10^−3^ in PBS and plated on to LB agar. The original stock was also diluted down 10^−8^ and plated. Plates were incubated overnight at 37°C and the number of colonies was counted and the values were adjusted to CFUs/ml for 10^0^. The data were plotted as a log decrease in colonies relative to the strain stock inoculum.

### Infection of HeLa cells

HeLa cells were taken from liquid nitrogen stocks and placed in a 75 ml tissue culture flask with 10 ml of complete DMEM media (DMEM supplemented with 2 mM L-glutamine, 10% (v/v) fetal bovine serum (FBS) and 1% (w/v) penicillin-streptomycin) and incubated at 37°C in 5% CO_2_. The next day the medium was removed and the cells were washed once with 10ml PBS and 10 ml of fresh complete DMEM medium was added. The cells were left for approximately 3 days until the cells were 80% confluent.

To prepare for infection assays, medium was removed from the confluent cells and the cells were washed once with 10 ml PBS. To detach the cells, 1 ml of trypsin was added and incubated for 5 mins at 37°C. After incubation, 10 ml of complete DMEM was added to the cells and pipetted up and down to remove clumps of cells. The cells were adjusted to a concentration of 1 × 10^5^ cells/ml and 1 ml of cells were added to each well of a 24 well plate to incubate overnight.

For infection, the *Salmonella* cultures were adjusted to a concentration of 5 × 10^6^ cells/ml in unsupplemented DMEM medium. The complete DMEM was removed from the HeLa cells and each well was washed twice with 1 ml of PBS. For each bacterial strain, 1 ml of bacterial cells (in plain DMEM) was added in triplicate to the wells containing HeLa cells. To initiate infection, the plates were centrifuged at 600 *g* for 5 min and left for 30 min at 37°C in 5% CO_2_ to allow infection. After 30 min the media was changed to complete DMEM with 100 μg/ml gentamycin to kill external bacteria. This was considered time 0 for invasion experiments. To prevent damage of the HeLa, the media was changed again to plain DMEM with 10µg/ml gentamicin after 1 hour of invasion. To end the experiment after 0 or 2 hours of invasion, the cells were washed twice with 1 ml of PBS. To lyse the HeLa cells, 1 ml of PBS with 0.5% Triton-X 100 was added to each well. The lysed cells were diluted down to 10^−3^ and plated and incubated overnight at 37°C. The following day the plates were counted and the CFUs/ml of the 10^0^ stocks were calculated.

### Infection of bone marrow derived macrophages

L-cell media is used to differentiate murine bone marrow cells to bone marrow derived macrophages. To prepare L-cell media, L-cells were thawed and added to 25 ml of complete RPMI (cRPMI) media (RPMI supplemented with 10% (v/v) FBS and 1% (w/v) penicillin-streptomycin) and transferred to a 75 ml tissue culture flask. The following day, the media was removed and dead cells were washed off with 10 ml of cRPMI, and 25 ml of fresh cRPMI was added and the cells were incubated at 37°C until confluent (approx. 2 days). Once confluent, the medium was removed and the cells were washed with 10 ml of PBS. The cells were detached from the flask by the addition of 2 ml of trypsin and incubated for 5 min at 37°C with gentle agitation. The cells were mixed in 10 ml and a total of 2.5 × 10^5^ cells were seeded into a 175 ml tissue culture flask with 50 ml of cRPMI. The cells were incubated until confluent (approx. 5 days). Once confluent, the media was removed and filtered through a 0.22 µM filter.

To obtain bone marrow derived macrophages, the bone marrow was isolated from mice aged 6-8 weeks. The bones of the hind legs were removed and placed into a petri dish with 5 ml of cRPMI. All excess muscle and skin was removed, the ends of each bone were cut and the bone marrow was rinsed through using cRPMI in a needle and syringe. The isolated bone marrow was centrifuged at 600 *g* for 10 min at room temperature. The pellet was re-suspended in 1 ml of red blood cell lysis buffer and incubated at room temperature for 5 min. 10 ml of cRPMI was added and the white blood cells were pelleted again and re-suspended in 10 ml of cRPMI and passed through a cell strainer into a fresh 50 ml tube. 5 × 10^6^ cells were seeded into a petri dish with 10 ml of cRPMI and 10% (v/v) L-cell medium. The cells were incubated overnight at 37°C in 5% CO_2_. The following day an additional 10 ml of cRPMI and 10% (v/v) L-cell medium was added to each petri dish. After 3 days, 10 ml of media was removed and replaced with 10 ml of fresh cRPMI and 10% (v/v) L-cell medium. After a further 3 days the cells were ready the invasion experiments.

For invasion experiments, macrophages were scraped from the bottom of the petri dish, centrifuged at 600 *g* for 10 min at 37°C and re-suspended in 5 ml of cRPMI. The cells were counted and diluted to a concentration of 1 × 10^5^/ml using cRPMI. For each invasion experiment 1 ml were placed in triplicate into a 12-well tissue culture plate and left overnight at 37°C in 5% CO_2_. The following day, bacterial cells were adjusted to a concentration of 1 × 10^6^ cells/ml in plain RPMI medium. For infection, the cRPMI was removed from the macrophages and each well was washed twice with PBS. For each bacterial strain, 1 ml of bacterial cells (in plain RPMI) was added in triplicate to the wells containing macrophages. To initiate infection, the plates were centrifuged at 600 *g* for 5 min at 37°C and incubated for 30 min at 37°C in 5% CO_2_. After 30 min the media was changed to plain RPMI with 100 µg/ml gentamycin to kill external bacteria. This was considered time 0 for invasion experiments. After 1 hour the media was changed to plain RPMI with 10 µg/ml gentamicin. To end the experiment after 2, 5 or 24 h of invasion, the cells were washed twice with PBS and mammalian cells were lysed in 1 ml of PBS with 1% (w/v) Triton-X 100 and were diluted down to 10^−3^. Dilutions were plated onto LB agar plates and incubated overnight at 37°C.

## Acknowledgments

This work was supported by a BBSRC grant to IRH. GLI was funded by the MIBTP BBSRC PhD Scholarship.

## Author contributions

GLI conducted the experimental work with help from JLR, AER, CAA, ES, TJM and CI. GLI wrote the manuscript with contributions from JAC and IRH. GLI lead the project with supervision from JAC and AFC, and IRH who conceived the study.

---

**Supplementary figure 1.**
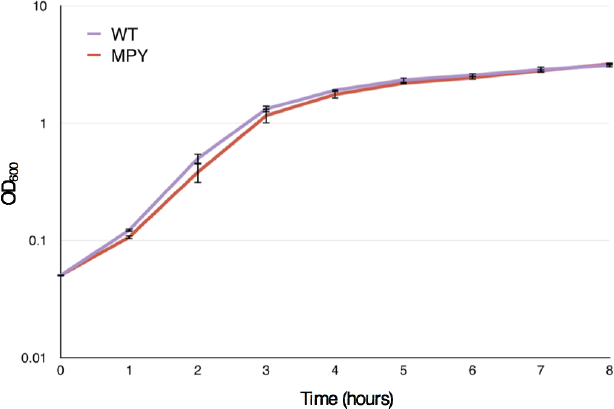
Growth of *S*. Typhimurium WT and MPY mutant in LB broth over 8 hours. Each point is the mean of 3 biological replicates and the error bars represent the standard deviation.

**Supplementary figure 2.**
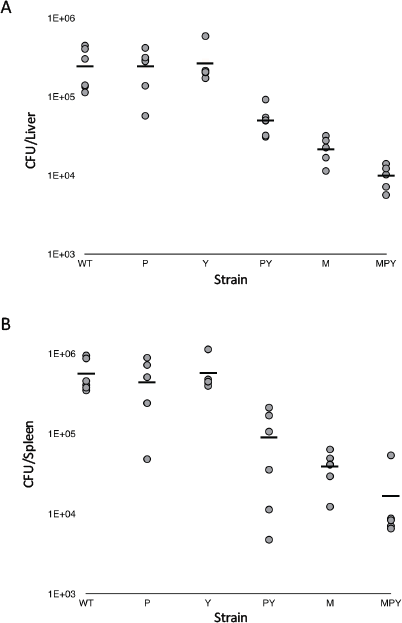
Bacterial burden of *Salmonella* Typhimurium WT and the *mlaD, pqiAB, yebST, pqiAByebST* and triple (MPY) mutants in C57BL/6 mice. Six mice were infected with the strains for 3 days via intraperitoneal injection. The CFUs/organ were counted in the liver **(A)** and spleen **(B)**. Each point represents the number of CFUs/organ from a single mouse, and the line indicates the mean number of CFUs. Mann-Whitney U test values were corrected for multiple testing for liver: WT vs P = 0.9, WT vs Y = 0.8, WT vs PY = 0.005, WT vs M = 0.005, WT vs MPY = 0.0065 and spleen: WT vs P = 0.6, WT vs Y = 0.7, WT vs PY = 0.013, WT vs M = 0.013, WT vs MPY = 0.013.

**Supplementary figure 3.**
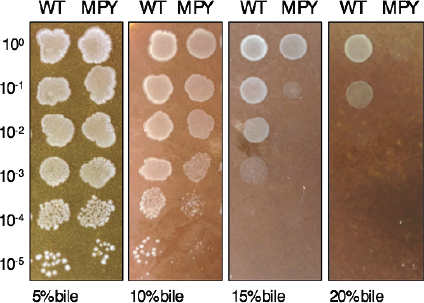
Bile sensitivity of MPY mutant. Logarithmic dilutions of the WT and MPY were tested on LB agar with increasing concentrations of bile.

**Supplementary figure 4.**
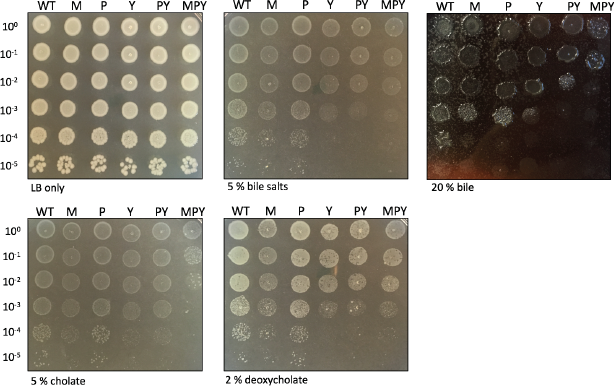
Sensitivity of MCE mutants on bile and components of bile. Logarithmic dilutions of the WT and *mlaD, pqiAB, yebST, pqiAByebST* and triple MCE mutant (MPY) on LB, bile salts, bile, cholate and deoxycholate.

